# Inherited long telomeres induce a genome-wide transcriptional response in budding yeast

**DOI:** 10.64898/2026.04.15.718807

**Authors:** Vasilisa Sidarava, David Lydall

**Affiliations:** Biosciences Institute, The Medical School, Framlington Place, Newcastle University, Newcastle upon Tyne, Tyne and Wear, NE2 4HH, UK

**Author notes:** Address correspondence to; Biosciences Institute, The Medical School, Framlington Place, Newcastle University, Newcastle upon Tyne, Tyne and Wear, NE2 4HH, UK.

**Keywords:** Telomere length, transcriptional response, *Saccharomyces cerevisiae*

## Abstract

Eukaryotes typically maintain telomere length within a defined range. While short telomeres are known to activate DNA damage responses and limit cell proliferation, long telomeres are associated with extended proliferative capacity. The broader cellular consequences of long telomeres are comparatively less well understood. In budding yeast *Saccharomyces cerevisiae*, long telomeres have been shown to influence gene expression at specific loci, but whether long telomeres affect transcription genome-wide has not been reported. Here, we analysed transcriptomes in a lineage that inherited long telomeres (originally due to a *rif2*Δ mutation). Transcriptomes were assessed over two rounds of mitosis and meiosis in the absence of the *rif2*Δ mutation. We show that strains with long telomeres exhibit a distinct gene expression profile, including upregulation of membrane transporters and downregulation of a smaller subset of genes. Both up- and down-regulated genes were distributed across the genome, arguing against a purely telomere-proximal effect on gene expression. Affected genes were enriched for Rap1 binding sites, consistent with a model in which long telomeres sequester telomere-associated transcriptional regulators, such as Rap1, and thereby affect gene expression at non-telomeric binding sites for these regulators. Accordingly, the magnitude of transcriptional changes was greatest in strains with the longest telomeres. Together, our findings demonstrate that long telomeres induce a genome-wide transcriptional response that can accompany inherited long telomeres across generations. Similar effects of long telomeres are likely to occur in other eukaryotes, including humans, where long telomeres are associated with disease.

**Article summary:** Telomeres protect chromosome ends, and their length is tightly regulated. While short telomeres are known to be harmful, the effects of long telomeres are less well understood. Using budding yeast, we show that inherited long telomeres alter the expression of dozens of genes across the genome, particularly membrane transporters. These changes are consistent with a model in which long telomeres sequester regulatory proteins away from other loci. Our findings may have broader implications in more complex organisms, including humans.

## Introduction

Telomeres are specialized nucleoprotein structures that cap the ends of linear chromosomes and protect the DNA from inappropriate repair, recombination, and degradation (Celli et al. 2006; Wu et al. 2006). Telomere length is typically maintained within an optimal range, and deviations from this range can have significant biological consequences (Armanios 2022).

In humans, abnormal telomere length is associated with distinct pathological syndromes. Short telomere syndromes are associated with impaired stem cell renewal and premature tissue degeneration, leading to manifestations such as bone marrow failure, pulmonary fibrosis, and liver disease (Armanios and Blackburn 2012). Conversely, long telomeres are associated with extended cellular proliferative potential and an increased risk of malignant transformation, likely by permitting additional rounds of cell division and thereby increasing the accumulation of mutations (McNally et al. 2019). At the cellular level, critically short telomeres activate DNA damage signalling pathways that can induce cell cycle arrest, senescence, or apoptosis (d’Adda di Fagagna et al. 2003). However, the effects of long telomeres beyond replicative lifespan remain unclear.

One challenge posed by excessively long telomeres is DNA replication, as telomeric DNA is intrinsically difficult to replicate due to its repetitive, G-rich sequence and associated protein complexes (Olson and Wuttke 2024). Consistent with this, long telomeres are unstable and can undergo recombination-based trimming events in yeast (Li and Lustig 1996), plants (Watson and Shippen 2007), and human cells (Pickett et al. 2009). Nevertheless, studies in budding yeast have not detected substantial defects in growth, lifespan, or genotoxic stress sensitivity in cells with long telomeres (Harari et al. 2017), suggesting that any detrimental effects are minimal.

In budding yeast, long telomeres can influence gene expression through at least two mechanisms. One is the telomere position effect (TPE), whereby telomere-associated silencing factors such as Rap1 and the Sir proteins promote transcriptional repression of genes located near chromosome ends through the spreading of silent chromatin (Gottschling et al. 1990; Brothers and Rine 2022). The other is by competing for DNA-binding regulatory proteins (Buck and Shore 1995; Marcand et al. 1996).

The effects of long telomeres on TPE can be complex. For example, in yeast that carried a single elongated telomere, the long telomere was associated with enhanced silencing of adjacent reporter genes (Kyrion et al. 1993). In contrast, *rif1*Δ mutants, with mostly elongated telomeres, showed reduced silencing at specific sub-telomeric reporter genes (Juarez-Reyes et al. 2023), suggesting that the effect of telomere elongation on TPE depends on the context.

Rap1 and the Sir2/3/4 complex also regulate transcription at internal genomic sites (Rusche et al. 2003; Azad and Tomar 2016). Consistent with this, long telomeres enhanced silencing at telomeres and weakened silencing at internal Sir- and Rap1-dependent loci (Buck and Shore 1995). Whether long telomeres induce broader transcriptional changes has not been reported. Here, we address this question by performing transcriptomic analysis of yeast strains with long telomeres, generated through mutation or inherited in the absence of the mutation that stimulated telomere elongation. Our analyses reveal a distinct genome-wide transcriptional response correlating with the inheritance of a long VIR telomere in yeast.

## Materials and methods

### Yeast strains

Strains used in this study are listed in Table 1.

**Table 1.**
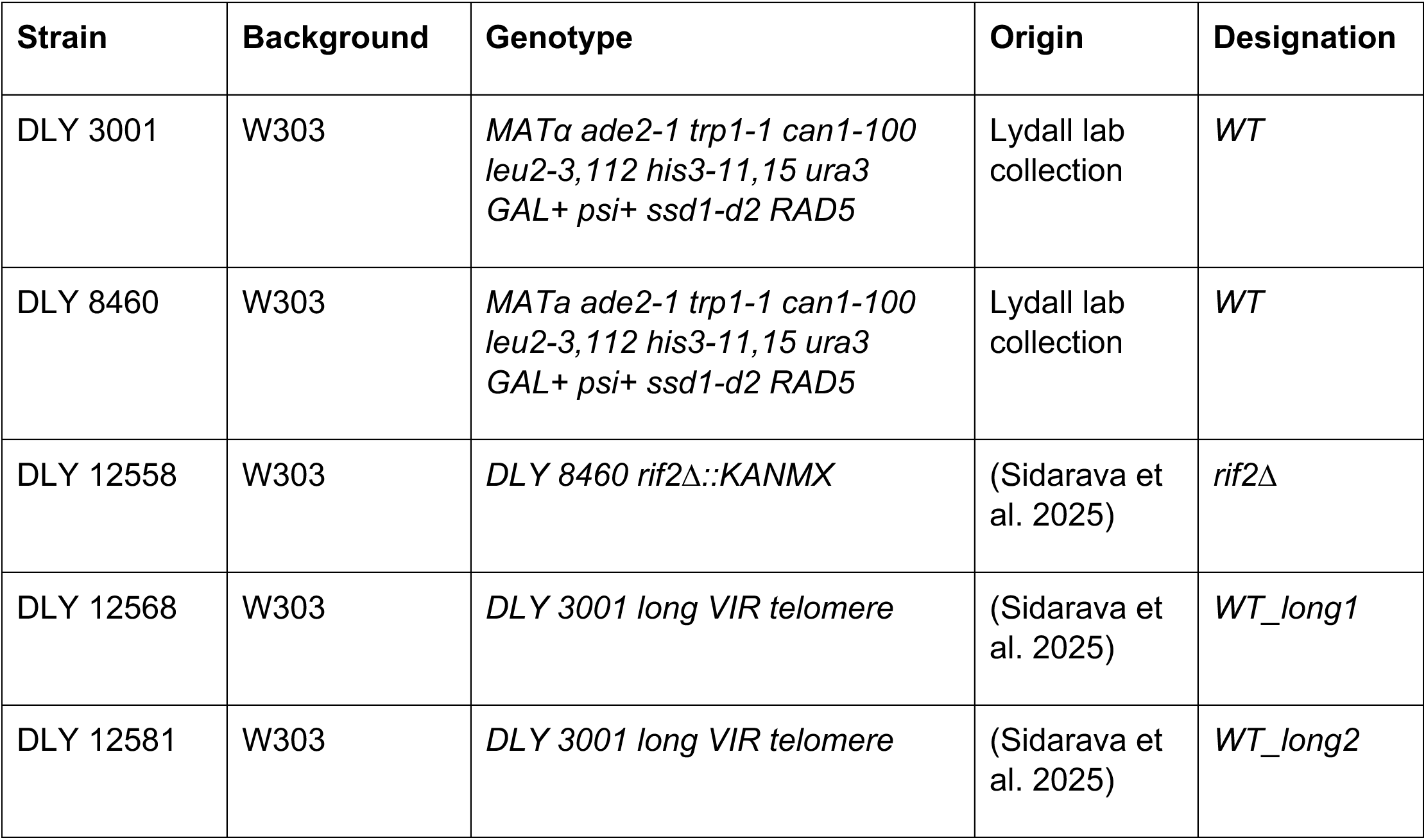
Strains used in this study.

### Yeast propagation and genetic manipulation

Yeast were routinely cultured at 30°C on YEPD plates (1% yeast extract, 2% peptone, 2% agar, 2% dextrose) supplemented with adenine (75 mg/l). To produce diploids, haploid strains of opposite mating types were mixed on YEPD plates and mated. Zygotes were picked after 3 hours at 30°C and isolated using a tetrad dissection microscope. Colonies derived from individual zygotes were grown on YEPD plates for 3.5 days and subsequently inoculated into 4 ml liquid YEPD and grown overnight at 30 °C. Each culture was then split into two equal aliquots: one used for genomic DNA extraction and telomere length analysis, and the other for RNA extraction and sequencing.

### Telomere length detection

Genomic DNA extraction and telomere length analysis were performed as previously described (Sidarava et al. 2025).

### RNA sequencing and downstream analyses

Total RNA was extracted using RNeasy Mini Kit (Qiagen) according to the manufacturer’s instructions. RNA sequencing was performed by Novogene Europe. Strand-specific libraries were generated from poly(A)-selected mRNA using the New England Biolabs NEBNext Ultra Directional RNA Library Prep Kit. After fragmentation and cDNA synthesis (with dUTP incorporation for strand specificity), libraries were adapter-ligated, size-selected, and treated with USER enzyme (NEB). This was followed by PCR amplification. Final library quality was assessed using a Bioanalyzer. Sequencing was carried out on the Illumina platform, yielding paired-end reads of 150 bp in length, with approximately 20 million reads per sample.

The initial bioinformatics analyses, including read filtering, alignment, transcript assembly, gene quantification, and functional enrichment (GO and KEGG), were carried out by Novogene Europe. Raw sequencing reads were quality-filtered using fastp (Chen et al. 2018) to remove adapters and low-quality reads. Clean reads were aligned to the *Saccharomyces cerevisiae* W303 reference genome (JRIU00000000) using HISAT2 (v2.2.1) (Kim et al. 2019). Transcript assembly was performed using StringTie (v2.2.3) (Pertea et al. 2015), and gene-level expression was quantified using featureCounts (v2.0.6) (Liao et al. 2014). Expression values were normalized as FPKM (Fragments Per Kilobase of transcript per Million mapped reads) (Trapnell et al. 2010) to perform principal component and cluster analyses.

Differential expression analyses were performed on raw read counts using the DESeq2 tool (v1.42.0) (Love et al. 2014). P-values were adjusted using the Benjamini–Hochberg false discovery rate (FDR) method. Genes with an adjusted p-value (padj) ≤ 0.05 were considered significantly differentially expressed.

Gene Ontology (GO) enrichment analysis was performed using the clusterProfiler (v4.8.1) R package (Wu et al. 2021), correcting for gene length bias. Enriched GO terms and pathways were considered significant at padj ≤ 0.05.

Heatmaps, Venn diagrams, and Gene Ontology enrichment analyses were generated using the NovoMagic platform (https://www.novogene.com/eu-en/novomagic-online-rna-seq-bioinformatics-analysis-tool/).

Spearman’s rank correlation was used to assess the relationship between fold-change magnitude and gene distance from the nearest telomere. Enrichment of Rap1 binding sites among shared DEGs was assessed using the hypergeometric test, with the total gene set defined as all genes detected by RNA-seq (n = 5696). Rap1 DNA-binding targets were obtained from the YEASTRACT database (Teixeira et al. 2023). The same test was used to assess overlap with genes affected by Rap1 depletion (Kalra et al. 2023), using their total gene set (n = 5088). Paired differences in fold-change magnitude were assessed using the Wilcoxon signed-rank test.

## Results

To test whether long telomeres influence gene expression, RNA sequencing was performed on diploid yeast strains that had different telomere lengths and were derived from a common lineage. Four diploid genotypes were analysed: a wild-type control (*WT/WT*), a heterozygous diploid carrying a mutation in the telomere length regulator *RIF2* which resulted in telomere elongation (*WT/rif2*Δ), and two genetically wild-type diploids with long VIR telomeres (*WT/WT_long1* and *WT/WT_long2*). All four diploids were generated by crossing haploid strains to the same wild-type haploid strain (Figure 1A). The *WT_long1* and *WT_long2* haploids were derived in a previous study through successive rounds of mating and meiosis that transmitted long VIR telomeres from a *rif2*Δ ancestor (Sidarava et al. 2025) (Figure 1A). As a result, *WT/rif2*Δ, *WT/WT_long1*, and *WT/WT_long2* are from a single lineage in which long telomeres were originally caused by a *rif2*Δ mutation.

**Figure 1.**
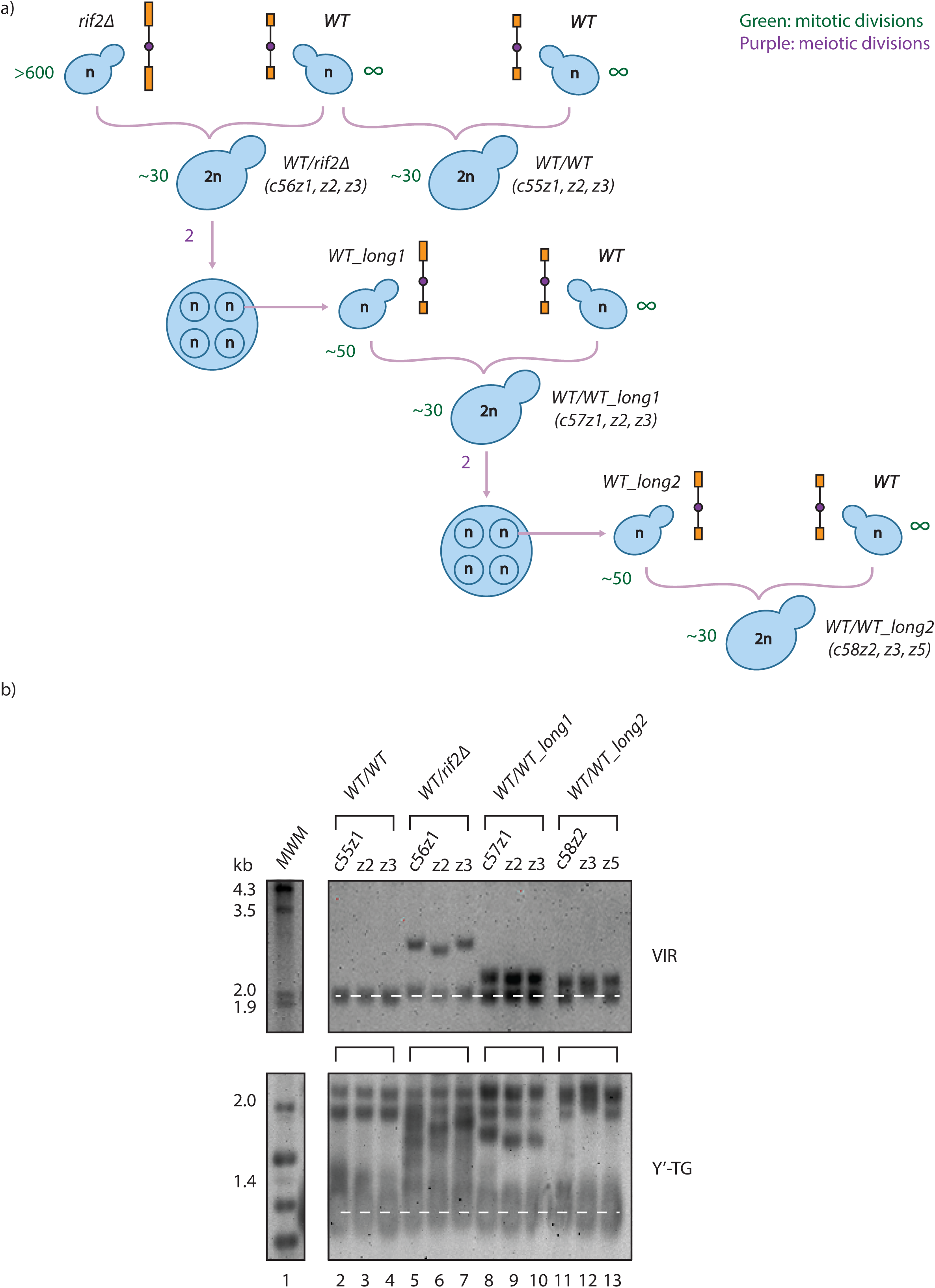
Origin and telomere length of strains used for transcriptome analysis. A) Lineage of strains analysed. A *rif2*Δ haploid was crossed to a wild-type haploid strain to generate a *WT/rif2*Δ diploid, which was sporulated to produce meiotic progeny. From this a wild-type haploid carrying long VIR telomeres (*WT_long1*) was isolated. *WT_long1* was crossed to the wild-type strain to generate a *WT/WT_long1* diploid, which was subsequently sporulated to yield a second haploid with long VIR telomeres (*WT_long2*). *WT_long2* was crossed to the wild-type strain to generate the *WT/WT_long2* diploid. Biological replicates for each diploid genotype examined are indicated in parentheses. Estimated numbers of mitotic and meiotic divisions undergone by haploid and diploid cells are indicated in green and purple, respectively, and were determined as described previously (Sidarava et al. 2025). The cells went through 4 further divisions in liquid culture prior to DNA extraction. B) Telomeric Southern blots. Three biological replicates for each genotype are shown. The *WT/WT* diploids served as controls. Telomere length was measured using VIR and Y’-TG probes. Dashed white line indicates normal telomere length.

Diploids were chosen for transcriptome analysis rather than haploids to minimise potential confounding effects of recessive background mutations that might be expressed in haploid cells (Szafraniec et al. 2003). In addition, diploid cells more closely reflect the chromosomal state of human somatic cells.

For each class of diploid, three biological replicates were obtained by isolating three independent zygotes from each cross. Diploid clones were expanded from zygotes, on plates and in liquid before cultures were split: half was used for DNA extraction and telomere length analysis, and the other half for RNA extraction and transcriptome analysis. This approach ensured that the transcriptome measured from each culture could be directly matched to telomere length measured from the same culture.

*WT/WT* strains had normal VIR telomere lengths – around 2.0 kb, as expected (Figure 1B). Consistent with our earlier work, two sets of VIR telomeres could be detected in *WT/rif2*Δ, *WT/WT_long1*, and *WT/WT_long2* strains, the shorter ones presumably derived from the wild-type parent and the longer ones from long-telomere parents (Sidarava et al. 2025). *WT/rif2*Δ cells had the longest long VIR telomeres (∼3.1 kb), while in *WT/WT_long1* and *WT/WT_long2* strains, the long VIR telomeres were 2.4 and 2.3 kb long, respectively. We estimate that there were 166 cell divisions between the segregation of the *rif2*Δ mutation and a long VIR telomere, and the cells in the *WT/WT_long2* cultures used for RNA extraction (Figure 1A). During these divisions VIR telomeres shortened gradually, most likely due to the end replication problem (Sidarava et al. 2025).

A Y′-TG probe was also used to examine average length of Y′ telomeres. Consistent with what we previously reported (Sidarava et al. 2025), the Y’-TG probe showed that *WT/rif2*Δ diploid cells had long Y’-telomeres (due to haploinsufficiency, or because it takes time for long telomeres to normalise in length). The *WT/WT*, *WT/WT_long1*, and *WT/WT_long2* cells had essentially normal length Y’-telomeres, but the *WT/WT_long1*, and *WT/WT_long2* cells had at least one long telomere (VIR). Overall, our Southern blots confirmed that the cultures we had harvested for transcriptome profiling contained the expected telomere lengths (of VIR and Y’).

To determine whether strains with long VIR telomeres shared transcriptional profiles, RNA sequencing data were analysed. First, principal component analysis (PCA) was performed to assess variation in global gene expression and the similarity between samples across all classes of diploids (Figure S1). Expression values were normalized using FPKM (Fragments per kilobase of transcript per million mapped reads), which accounts for transcript length and sequencing depth. Reassuringly, most biological replicates clustered together, indicating consistency of gene expression profile with groups of clones. However, sample c56z1 (*WT/rif2*Δ) showed strong deviation from its group, separating clearly from all other samples along the first principal component. Standard sequencing quality metrics, including clean read percentage, GC content distribution, error rate, and mapping rate, were comparable in c56z1 and the other samples, suggesting that the deviation reflects biological rather than technical variability. Because the inclusion of this sample disproportionately influenced the variance structure used for group comparisons, it was excluded from the primary analyses presented here. All analyses that follow were also repeated with c56z1 included (File S2), and similar results obtained.

To identify transcriptional changes associated with long telomeres, pairwise differential expression analyses were performed between each long-telomere strain and the *WT/WT* control using DESeq2 (see Methods). For each gene, fold changes and adjusted p-values were calculated. Lists of significantly differentially expressed genes (DEGs; adjusted p ≤ 0.05) for all three comparisons are provided in File S1.

Inspection of individual DEGs by pairwise comparison revealed upregulation of diverse transporter genes including phosphate, amino acid, and sugar transporters (File S1). In addition, the starvation marker *YGP1* was upregulated (Destruelle et al. 1994), while ribosomal and mitochondrial genes were downregulated. This pattern is consistent with a starvation-like transcriptional response (Wu et al. 2004; Petti et al. 2011). Histone genes from all four core histone families (H2A, H2B, H3, and H4) were downregulated in strains that inherited long telomeres (*WT/WT_long1* and *WT/WT_long2*; fold change 0.63–0.85x; File S1), as also seen under nutrient limitation (Chatzitheodoridou et al. 2024).

To compare transcriptional patterns across strains, all significant DEGs (adjusted p-value ≤ 0.05) from the three pairwise comparisons were combined and subjected to hierarchical clustering. The resulting heatmap showed clear separation between *WT/WT* samples and all long-telomere strains, indicating that long telomeres induce a distinct transcriptional profile (Figure 2). Replicates of *WT/WT_long1* and *WT/WT_long2* generally clustered together, supporting the reproducibility of inherited long-telomere effects across generations. One *WT/WT_long2* replicate clustered closer to *WT/rif2*Δ in the heatmap; however, this variation was not evident in the PCA, where all *WT/WT_long2* replicates clustered tightly (Figure S1).

**Figure 2.**
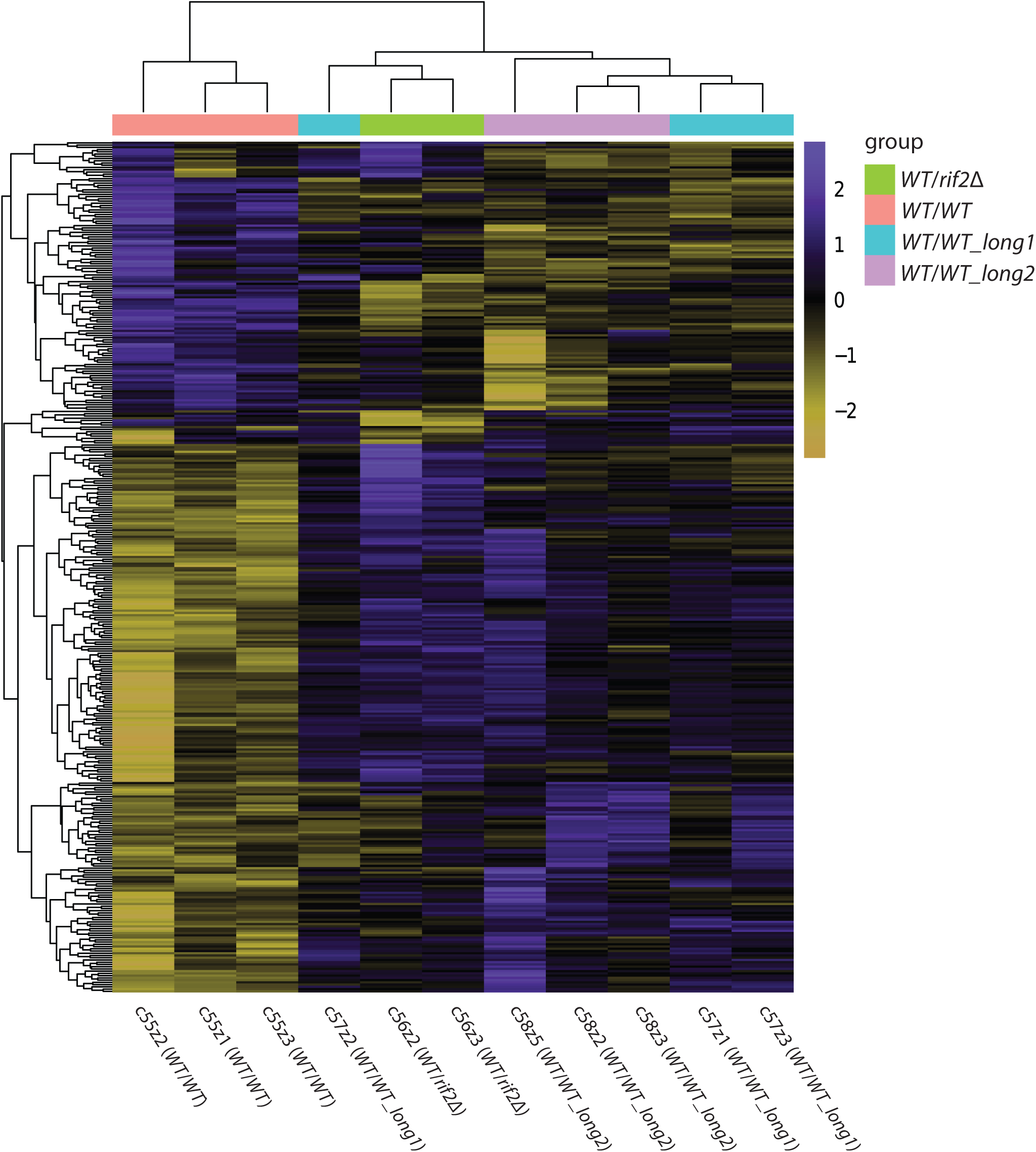
Long telomere strains (where long telomeres are either inherited or mutation-induced) share a distinct transcriptional profile from wild type. Heatmap of combined significantly differentially expressed genes (DEGs; adjusted p ≤ 0.05) from pairwise comparisons between long telomere strains and the wild type. Each column represents a sample (replicate), and each row represents a gene. Expression values were normalized as FPKM. For visualization, values were also row-normalized using Z-scores (colour intensity reflects each gene’s expression relative to its mean across all samples). Blue colour indicates upregulation and yellow colour indicates downregulation.

The overlap between DEGs identified in the three pairwise comparisons was visualized using a Venn diagram (Figure 3). Fifty-seven DEGs were shared across all long-telomere conditions, defining a common transcriptional response associated with long telomeres (Table 2). In addition, each comparison contained a substantial number of unique DEGs (for example, 103 genes unique to *WT/WT_long2* vs *WT/WT*), suggesting that long telomeres might also produce strain- or generation-specific transcriptional effects.

**Figure 3.**
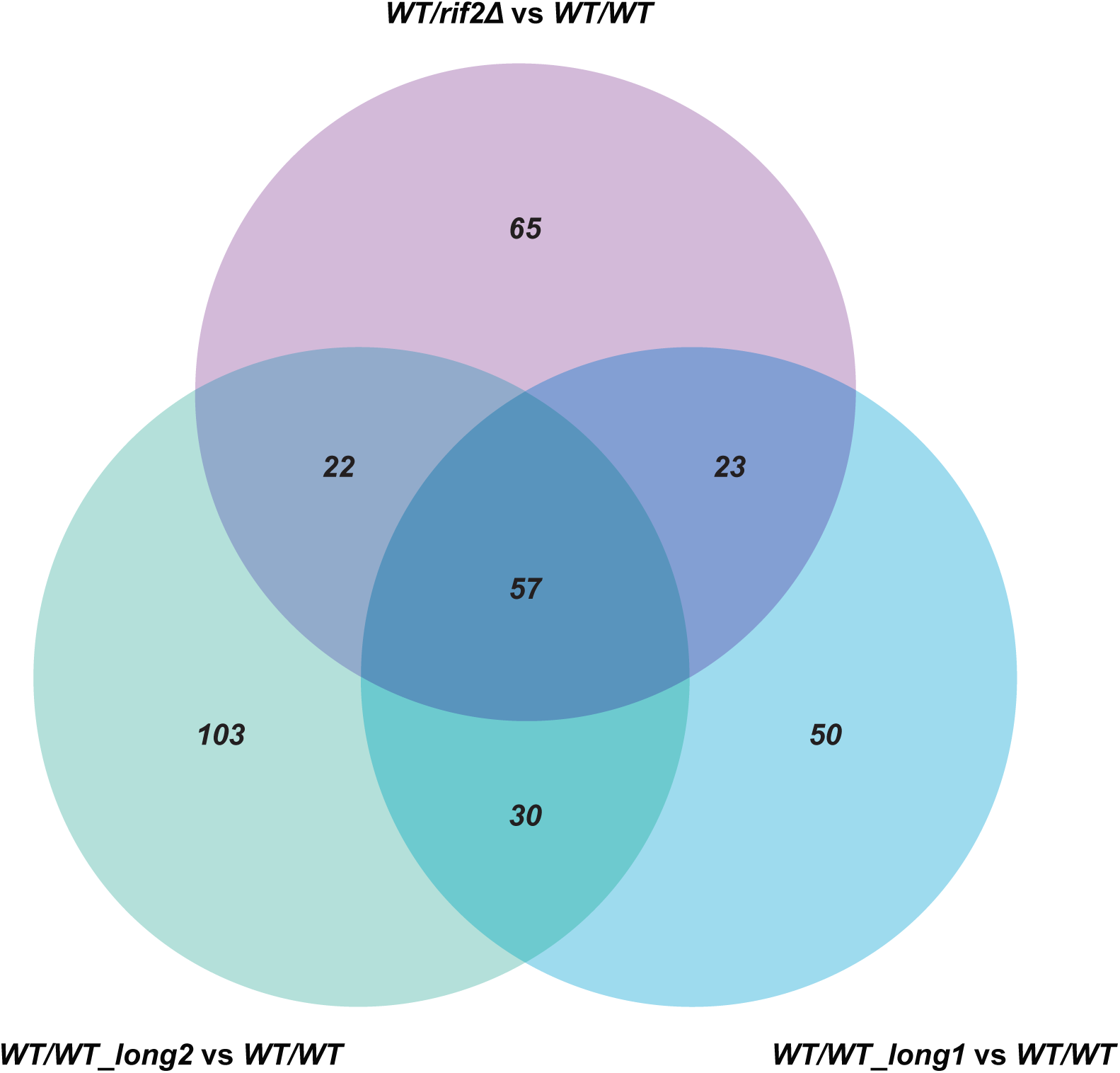
Long-telomere strains share a core set of differentially expressed genes. Venn diagram illustrating the overlap between significantly differentially expressed genes (adjusted p ≤ 0.05) from three pairwise comparisons: *WT/WT_long1* vs *WT/WT* (blue), *WT/WT_long2* vs *WT/WT* (green), and *WT/rif2*Δ vs *WT/WT* (purple). Numbers in the diagram indicate DEGs unique to a given comparison (non-overlapping regions) or shared between comparisons (overlapping regions).

**Table 2.** Genomic position and Rap1-binding status of the 57 DEGs.

| Gene ID | Gene standard name and function | Chromosome number | Distance from telomere (kb) | Binds Rap1? |
| --- | --- | --- | --- | --- |
| *YML128C* | *MSC1* – Meiotic sister chromatid recombination protein 1 | XIII | 15 | No |
| *YDR536W* | *STL1* – Sugar transporter | IV | 23 | No |
| *YML123C* | *PHO84* – Inorganic phosphate transporter | XIII | 24 | Yes |
| *YKL216W* | *URA1* – Dihydroorotate dehydrogenase | XI | 25 | No |
| *YNR067C* | *DSE4* – Glucan endo-1,3-beta-D-glucosidase involved in cell separation after cytokinesis | XIV | 29 | No |
| *YDL237W* | *AIM6* – Altered inheritance of mitochondria protein 6 | IV | 30 | No |
| *YLL055W* | *YCT1* – High affinity cysteine transporter | XII | 30 | Yes |
| *YBR286W* | *APE3* – Vacuolar aminopeptidase Y | II | 39 | No |
| *YPL265W* | *DIP5* – Dicarboxylic amino acid permease | XVI | 41 | No |
| *YDL223C* | *HBT1* | IV | 57 | No |
| *YHR202W* | *SMN1* – Putative AMP hydrolase | VIII | 60 | No |
| *YHL016C* | *DUR3* – Urea active transporter | VIII | 72 | Yes |
| YKL195W | *MIA40* – Import and assembly protein in mitochondrial intermembrane space | XI | 76 | No |
| YGR260W | *TNA1* – High-affinity nicotinic acid transporter | VII | 78 | Yes |
| YCR068W | *ATG15* – Putative lipase | III | 79 | Yes |
| YDL210W | *UGA4* – GABA-specific permease | IV | 84 | No |
| YKL187C | *FAT3* – Fatty acid transporter | XI | 89 | No |
| YCR061W | *YCV1* – Uncharacterized membrane protein YCR061W | III | 91 | No |
| YDL204W | *RTN2* – Reticulon-like protein 2 | IV | 94 | Yes |
| YOL119C | *MCH4* – Probable transporter | XV | 95 | Yes |
| YDR492W | *IZH1* – ADIPOR-like receptor | IV | 97 | No |
| YJL172W | *CBPS* – Carboxypeptidase S | X | 98 | No |
| YFL016C | *MDJ1* – DnaJ homolog 1, mitochondrial | VI | 104 | Yes |
| YPR158W | *CUR1* – Curing of [URE3] protein 1 | XVI | 105 | No |
| YDR483W | *KRE2* – Glycolipid 2-α-mannosyltransferase | IV | 110 | No |
| novel.119 | *TPO3* – Polyamine transporter 3 | XVI | 111 | No |
| YJL163C | *YJQ3* – Uncharacterized membrane protein YJL163C | X | 112 | No |
| YML075C | *HMDH1* – 3-hydroxy-3-methylglutaryl-coenzyme A reductase 1 | XIII | 115 | No |
| YMR266W | *RSN1* – Uncharacterized protein | XIII | 126 | Yes |
| YBR235W | *VHC1* – Vacuolar cation-chloride cotransporter 1 | II | 127 | No |
| YNL268W | *LYP1* – Lysine-specific permease | XIV | 138 | Yes |
| YCR017C | *CWH43* | III | 144 | No |
| YGL186C | *TPN1* – Vitamin B6 transporter | VII | 151 | No |
| YIL030C | *DOA10* – ERAD-associated E3 ubiquitin-protein ligase | IX | 154 | No |
| YCR021C | *HSP30* – 30 kDa heat shock protein | III | 160 | No |
| YNL237W | *YTP1* – Protein | XIV | 205 | No |
| YHR096C | *HXT5* – Probable glucose transporter | VIII | 268 | Yes |
| YNL195C | *YNT5* – Uncharacterized protein<br>YNL195C | XIV | 271 | No |
| YLR069C | *EFGM* – Elongation factor G, mitochondrial | XII | 271 | No |
| YKL034W | *TUL1* – Transmembrane E3 ubiquitin-protein ligase 1 | XI | 295 | No |
| YOR002W | *ALG6* – Dolichyl pyrophosphate Man9GlcNAc2 alpha-1,3-glucosyltransferase | XV | 329 | No |
| *YNL160W* | *YGP1* – Protein YGP1 | XIV | 336 | No |
| *YMR034C* | *RCH1* – Solute carrier | XIII | 339 | No |
| *YBR054W* | *YRO2* | II | 343 | No |
| *YLR299W* | *ECM38* – Glutathione hydrolase proenzyme | XII | 352 | Yes |
| *YNL142W* | *MEP2* – Ammonium transporter | XIV | 357 | Yes |
| *YDL054C* | *MCH1* – Probable transporter | IV | 360 | Yes |
| *YBR067C* | *TIP1* – Temperature shock-inducible protein 1 | II | 372 | No |
| *YBR072W* | *HSP26* – Small heat shock protein with chaperone activity | II | 382 | Yes |
| *YGL055W* | *OLE1* – Delta(9) fatty acid desaturase | VII | 399 | Yes |
| *YOR187W* | *EFTU* – Elongation factor Tu, mitochondrial | XV | 407 | No |
| *YGL008C* | *PMA1* – Plasma membrane ATPase 1 | VII | 479 | Yes |
| *YGR060W* | *ERG25* – C-4 methylsterol oxidase | VII | 480 | Yes |
| *YGL006W* | *ATC2* – Calcium-transporting ATPase 2 | VII | 486 | Yes |
| *YLR207W* | *HRD3* – ERAD-associated E3 ubiquitin-protein ligase component | XII | 522 | No |
| *YDR135C* | *YCF1* – Metal resistance protein | IV | 723 | Yes |
| *YDR205W* | *MSC2* – Probable zinc transporter | IV | 772 | No |

To explore whether the set of 57 DEGs represented specific functional categories, Gene Ontology enrichment analysis was performed. This revealed significant overrepresentation of transport-related categories (Figure S2), consistent with the transporter induction observed in the individual comparisons. No other functional categories showed strong enrichment.

Interestingly, subtelomeric regions are enriched in stress-responsive and transporter genes (Ai et al. 2002; Brown et al. 2010). Thus, we hypothesized that the change in expression of transporters and other shared DEGs could be explained by changes in subtelomeric chromatin induced by long telomeres. For example, long telomeres could affect telomere position effect (TPE), whereby the expression of genes in proximity to long telomeres is repressed. However, only 7 of the 57 shared DEGs were located within 30 kb of a chromosome end (Table 2), and no significant correlation was observed between the fold change and distance from telomeres in any strain (Figure 4). Together, these observations argue against telomere position effect as the primary driver of the shared transcriptional response.

**Figure 4.**
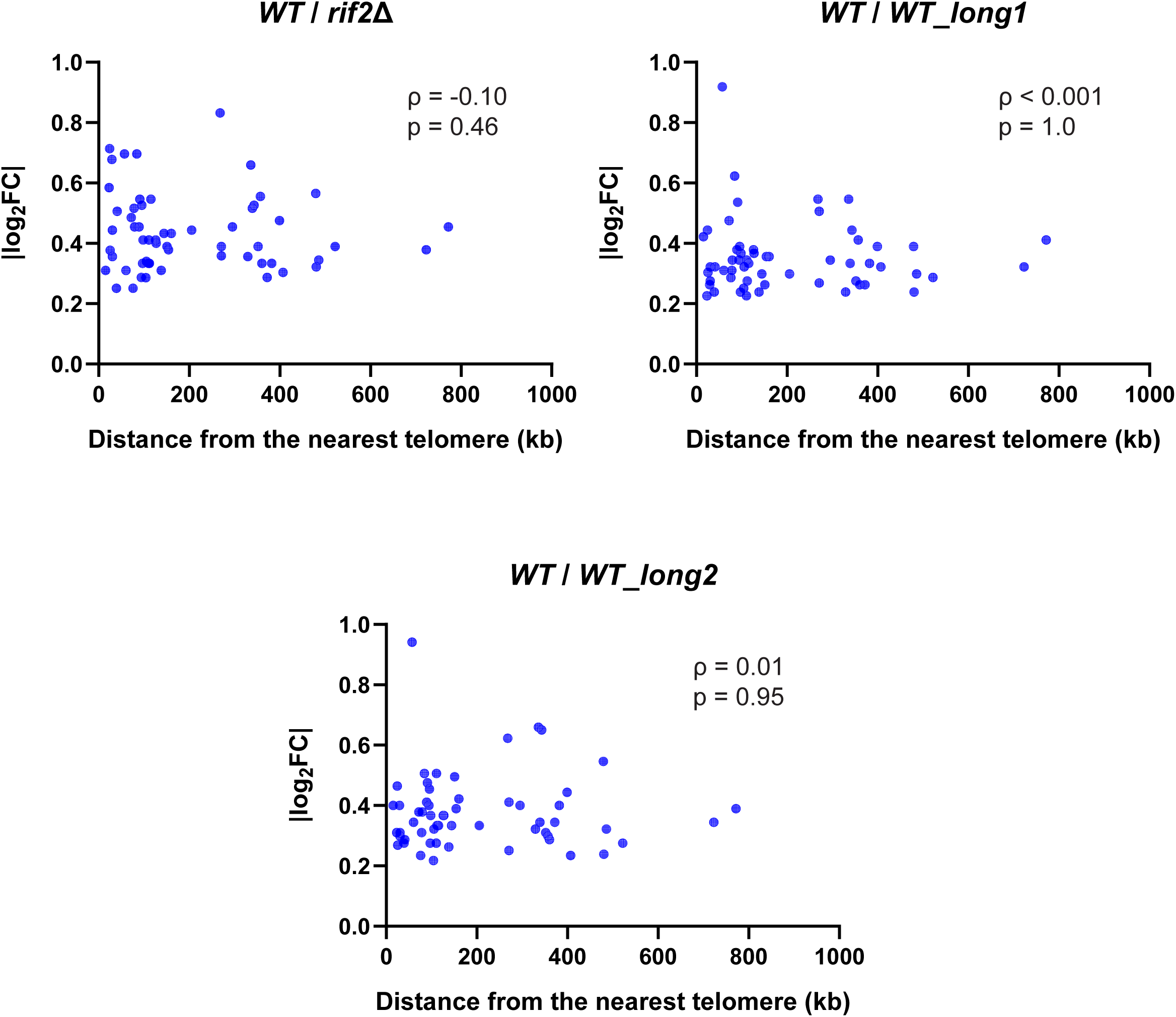
The observed transcriptional changes do not correlate with the distance from the nearest telomere. Scatter plots showing the relationship between fold-change magnitude (|log_2_FC|) and distance from the nearest telomere (kb) for the 57 DEGs in *WT/rif2*Δ, *WT/WT_long1*, and *WT/WT_long2* strains. Each point represents a gene. Spearman’s rank correlation was used to assess the relationship between variables, with the corresponding ρ and p values indicated on each plot.

One possible explanation for the observed genome-wide changes in gene expression is the sequestration of transcriptional regulators away from their internal genomic targets by telomeric DNA. Rap1 is a good candidate of such a regulator: in addition to binding telomeric DNA, Rap1 is a transcriptional regulator with 1582 documented DNA-binding targets across the genome (YEASTRACT (Yeast Search for Transcriptional Regulators And Consensus Tracking) database (Teixeira et al. 2023)). If an elongated telomere reduces Rap1 occupancy at non-telomeric promoters, this could lead to misregulation of those genes. To test this hypothesis, the 57 shared DEGs were cross-referenced with YEASTRACT annotations: 20 of the 57 showed documented Rap1 DNA-binding evidence (Table 2). This proportion is higher than expected by chance (hypergeometric test, p ≈ 0.02, based on 5696 genes detected by RNA-seq, of which 1285 bind Rap1), consistent with a model in which long telomeres alter Rap1 availability at some of its internal targets and thereby contribute to the shared transcriptional response. The Sir2/3/4 complex, which also functions at both telomeres and internal loci, is another candidate for such redistribution. However, none of the 57 shared DEGs overlapped with SIR-dependent genes identified at native telomeres (Ellahi et al. 2015), arguing against SIR-mediated silencing as a contributor to the observed response. In addition, because neither subtelomeric localization nor Rap1/SIR binding accounts for all shared DEGs, additional regulatory mechanisms or indirect downstream effects are likely to also contribute to the transcriptional consequences of telomere elongation.

Previous work has shown that partial depletion of Rap1 using a doxycycline-repressible system results in widespread transcriptional changes, with distinct gene sets responding at different levels of Rap1 abundance (Kalra et al. 2023). However, overlap between our 57 DEGs and genes responsive to Rap1 depletion was inconsistent across depletion levels (Table S1). At approximately 60%, 40%, and 20% of normal Rap1 levels, the Kalra et al. dataset identified 1221, 1920, and 2341 affected genes, respectively, of which 17, 26, and 24 overlapped with our 57 DEGs. Only the intermediate depletion level showed significant enrichment (hypergeometric test: p = 0.188, 0.022, and 0.324, respectively; total number of genes analyzed by Kalra et al. – 5088). This may reflect differences between partial Rap1 loss and partial redistribution of Rap1 predicted by our model. Direct tests of Rap1 occupancy at promoters in long-telomere strains would be needed to confirm redistribution.

Since telomere length decreased between *WT/rif2*Δ cells and *WT/WT_long1* cells and further in *WT/WT_long2* cells (Figure 1B), we hypothesised that the magnitude of changes in levels of DEGs would change accordingly. The changes for each gene were compared between pairs of strains (Figure 5). Fold changes were significantly reduced from *WT*/*rif2*Δ to *WT*/*WT_long1* (Wilcoxon signed-rank test, p < 0.0001; median Δ|log_2_FC| = 0.077) but did not decrease further between *WT*/*WT_long1* and *WT*/*WT_long2* (slight increase in magnitude, p = 0.0051; median Δ|log_2_FC| = 0.024). This pattern is broadly consistent with the telomere length data. In *WT/rif2*Δ diploids, the *rif2*Δ haploid parent contributed elongated telomeres at all 32 chromosome ends, so approximately half of all telomeres in the diploid are expected to be long, with VIR telomeres reaching ∼3.1 kb. In contrast, *WT/WT_long1* and *WT/WT_long2* diploids carried normal Y’-telomeres and presumably only a subset of long telomeres, including VIR, which were detectable but shorter than in the *WT/rif2*Δ diploid (∼2.4 and ∼2.3 kb, respectively) (Figure 1B). The large reduction in total long-telomere burden between *WT/rif2*Δ and *WT/WT_long1* likely contributes to the corresponding drop in transcriptional response, while the similar VIR lengths in *WT/WT_long1* and *WT/WT_long2* are consistent with their comparable expression profiles. These observations are consistent with a model in which the transcriptional response scales with the amount of Rap1 sequestered at elongated telomeric DNA.

**Figure 5.**
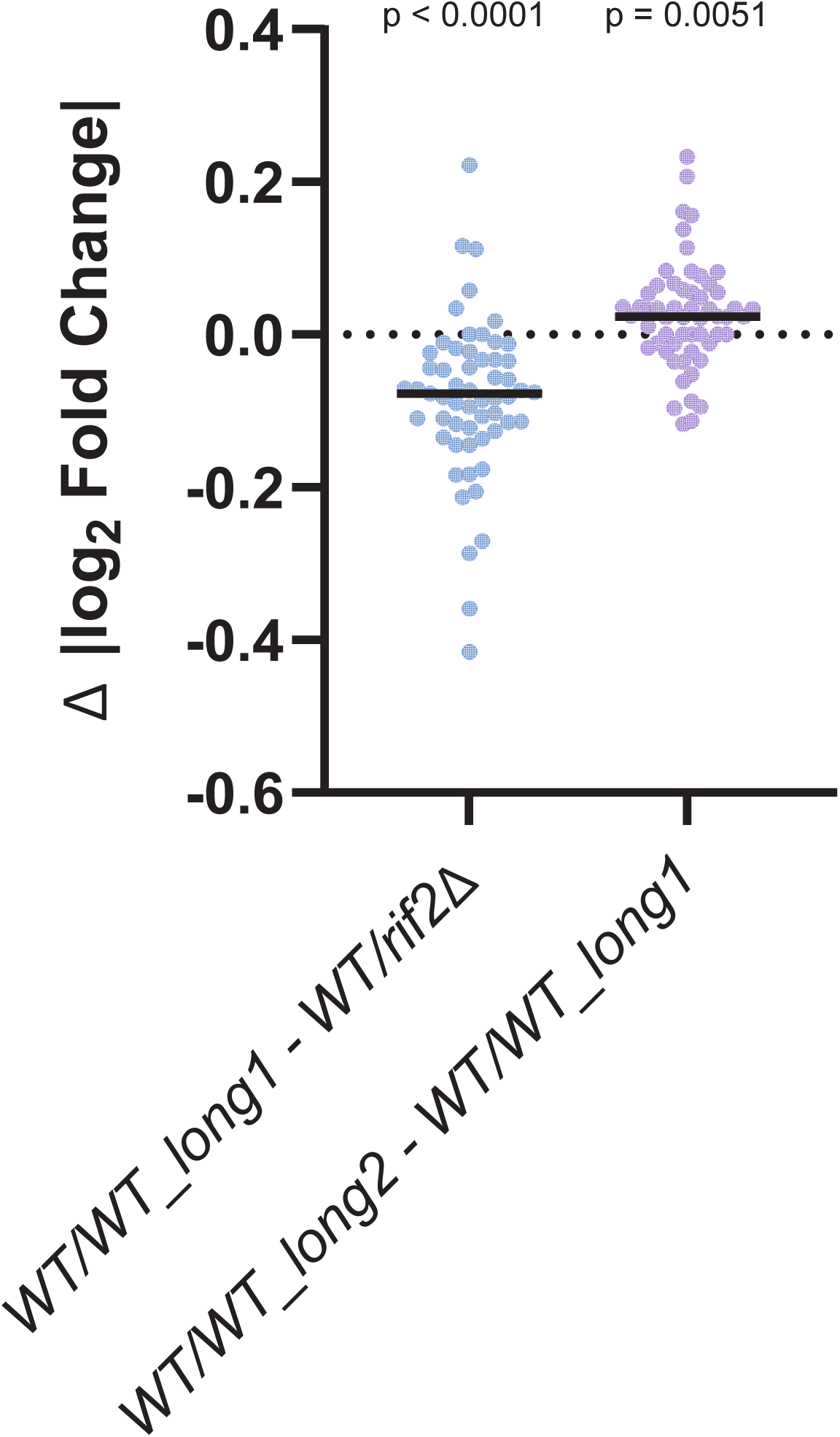
Per-gene changes in |log_2_FC| across long-telomere strains. Paired differences in absolute log_2_ fold-change (|log_2_FC|) values were calculated for each of the 57 shared DEGs between *WT/WT_long1* and *WT/rif2*Δ, and between *WT/WT_long2* and *WT/WT_long1*. Each point represents one gene. The dashed line indicates zero difference. Solid lines indicate medians. A Wilcoxon signed-rank test was used to assess whether paired |log_2_FC| differences were significantly different from zero; p-values are shown.

## Discussion

RNA-seq analysis of three generations of yeast strains, in which long telomeres were initially generated by a *rif2*Δ mutation and subsequently inherited by wild-type cells, revealed a genome-wide transcriptional response to long telomeres. A core set of 57 genes was consistently differentially expressed across all long-telomere strains. Upregulated genes were enriched for membrane transporters, while downregulated genes included several genes involved in mitochondrial translation and import. This pattern resembles transcriptional responses reported under nutrient deprivation (Wu et al. 2004).

We considered whether the shared transcriptional response could be explained by telomere position effect (subtelomeric gene silencing). Only a small fraction of the shared DEGs identified here were located within 30 kb of chromosome ends, and fold change magnitude did not correlate with distance from telomeres. This argues against altered subtelomeric silencing as the primary driver of the shared transcriptional response to telomere elongation. In contrast, the enrichment of Rap1 binding among shared DEGs supports a model in which long telomeres influence transcription through redistribution of Rap1. Rap1 functions at both telomeric and non-telomeric loci and can act as either an activator or a repressor of transcription. Increased sequestration of Rap1 at elongated telomeres could therefore reduce its availability at internal promoters, providing a potential explanation for both up- and down-regulated DEGs. Consistent with this model, the strongest changes in gene expression were observed in *WT/rif2*Δ strains, which had the longest telomeres, followed by smaller changes in *WT/WT_long1* and *WT/WT_long2*.

An alternative, non-exclusive explanation for the transcriptional response to long telomeres is that long telomeres impose a metabolic burden on the cell. Replication and maintenance of elongated telomeric DNA could increase demand for nucleotides or energy, leading to a physiological state that resembles nutrient limitation and triggers compensatory transporter induction. A link between nutrient sensing and telomere maintenance has been reported: inhibition of TOR signalling has been linked to telomere shortening, suggesting crosstalk between metabolic state and telomere regulation (Ungar et al. 2011; Kupiec and Weisman 2012). However, we did not detect changes in expression of TOR pathway components in our strains. Additionally, in the most extreme case (*WT/rif2*Δ), we estimate that elongated telomeres contribute approximately 35 kb (about 1.1 kb extra per telomere) of additional telomeric DNA, representing less than 0.15% of the diploid genome. Thus, it is unlikely that such a small increase in replication load is sufficient to alter cellular metabolism.

Since in our experiments the diploid wild-type strains with long telomeres were derived from a *WT/rif2*Δ diploid, we considered whether the transcriptional response might reflect long-lasting telomere-length-independent consequences of the *rif2*Δ mutation. However, the same core transcriptional signature (57 DEGs) was observed in wild-type segregants with long telomeres after meiosis and subsequent propagation, indicating that long telomeres, rather than the *rif2*Δ mutation itself, are the primary driver of the observed expression changes.

Overall, the magnitude of gene expression changes between long-telomere strains and wild-type controls was modest (typically <2-fold for the 57 DEGs). It is well established, however, that even modest changes in gene dosage can have significant phenotypic consequences. In human aneuploidies such as Down syndrome, a 1.5-fold increase in chromosome 21 gene copy number and presumably gene expression is sufficient to cause this complex multi-system disorder (Antonarakis 2017). Dosage-sensitive genes more broadly associate with disease phenotypes including cardiac defects, cancers, and neuropsychiatric disorders (Rice and McLysaght 2017). In *S. cerevisiae*, subtle shifts in gene expression can influence cellular behaviour under specific environmental conditions (Rest et al. 2013; Keren et al. 2016), or in combination with other mutations.

Telomere length-associated effects on gene expression have been reported in other eukaryotic systems. In human cells, telomere position effect has been documented both locally (Baur et al. 2001) and over long genomic distances (Kim and Shay 2018). At the same time, multiple telomeric proteins, including RAP1 and other shelterin components, have been shown to bind internal genomic sites and regulate transcription (Yang et al. 2011). However, whether telomere length modulates this regulation genome-wide has not been tested. Our findings demonstrate that long telomeres induce a genome-wide transcriptional response in yeast, with Rap1 targets enriched among the affected genes. Whether similar length-dependent transcriptional responses occur in mammalian cells, and whether they involve redistribution of telomere-associated factors, remains an important question for future study.

## Data availability

Raw RNA-seq reads and raw gene counts for all samples have been deposited in NCBI’s Gene Expression Omnibus (Edgar et al. 2002) and are accessible through GEO Series accession number GSE327262 (https://www.ncbi.nlm.nih.gov/geo/query/acc.cgi?acc=GSE327262). Differentially expressed gene lists are provided in Files S1 and S2. All strains used in this study are available upon request.

## Acknowledgements

We thank Ed Louis, Laura Maringele, and Simon Whitehall for helpful discussions and advice.

---

**Figure S1.** *Biological replicates clustered consistently by genotype, apart from a single outlier.* Principal component analysis (PCA) of RNA-seq samples based on log₂(FPKM + 1) values. Each point represents one biological replicate, colour- and shape-coded by genotype group. The first two principal components (PC1 and PC2) explain 25.73% and 15.90% of the total variance, respectively. Analyses were also performed including the outlier sample (File S2). Inclusion of this sample would not substantially alter the overall patterns of differential expression or the main conclusions of this study.

**Figure S2.** *Upregulation of transport-related genes represents one of the most consistent responses to telomere elongation.* A scatterplot showing enriched Gene Ontology (GO) terms for differentially expressed genes (DEGs). The x-axis indicates the gene ratio (the fraction of DEGs annotated to a given GO term relative to all DEGs with GO annotations). The y-axis lists GO terms. Dot size reflects the number of DEGs assigned to each term, and dot colour indicates enrichment significance (adjusted p-value).

**Table S1.** Overlap between the 57 DEGs and genes affected by Rap1 depletion (Kalra et al. 2023).

**File S1.** Lists of differentially expressed genes across the three pairwise comparisons, excluding the outlier RNA-seq replicate c56z1 (WT/rif2Δ).

**File S2.** Analyses including the outlier RNA-seq replicate c56z1 (WT/rif2Δ), comprising pairwise comparisons tables, cluster analysis, Venn diagram of overlapping DEGs, Gene Ontology analysis of overlapping DEGs, a table listing overlapping DEGs, analysis of correlation between |log_2_FC| and distance from the nearest telomere, and per-gene changes in |log_2_FC| across long-telomere strains.

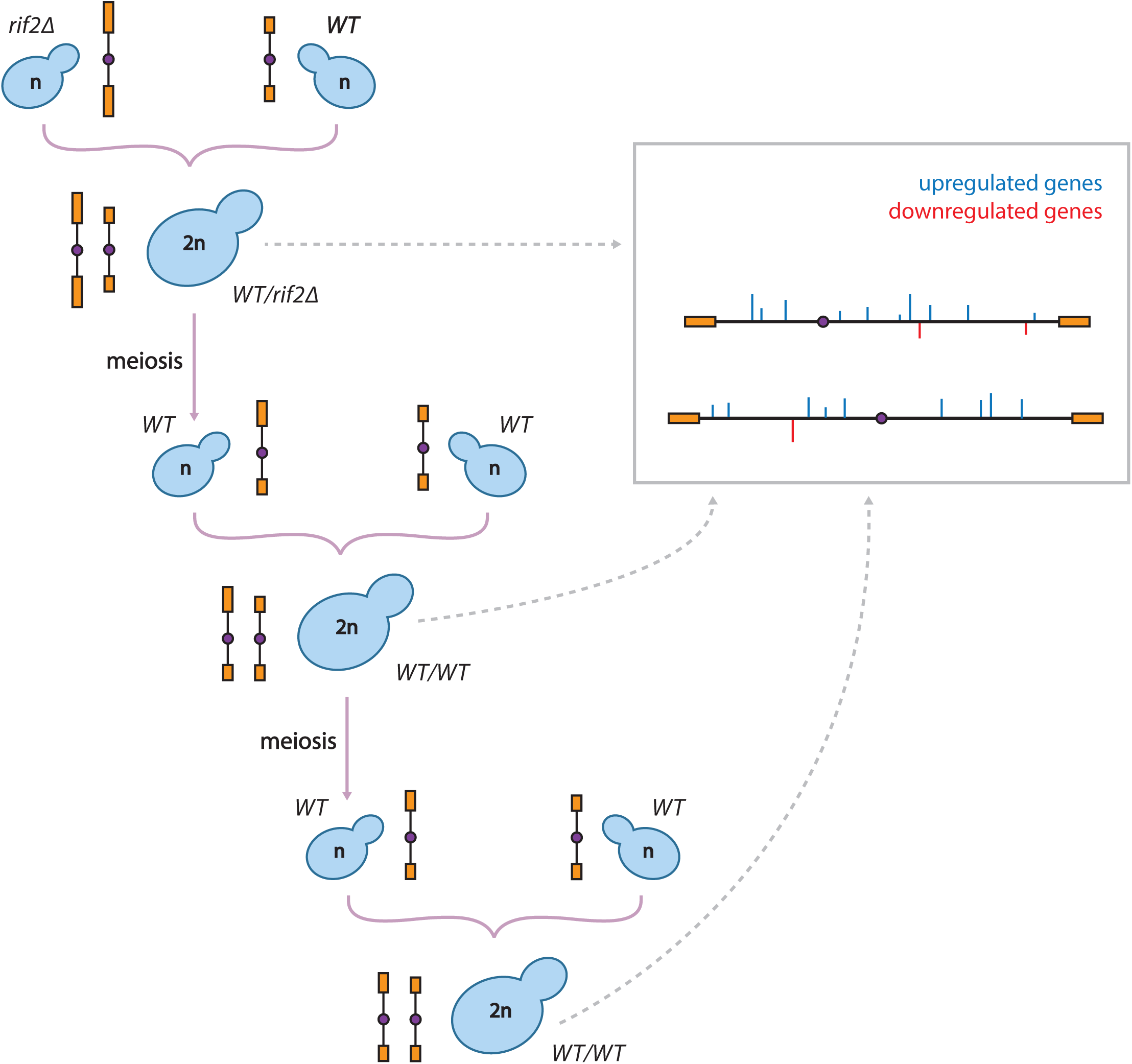

## References

Ai W et al. 2002. Regulation of subtelomeric silencing during stress response. Mol Cell. 10(6):1295–1305. doi: 10.1016/s1097-2765(02)00695-0.

Antonarakis SE. 2017. Down syndrome and the complexity of genome dosage imbalance. Nat Rev Genet. 18(3):147–163. doi: 10.1038/nrg.2016.154.

Armanios M, Blackburn EH. 2012. The telomere syndromes. Nat Rev Genet. 13(10):693–704. doi: 10.1038/nrg3246.

Armanios M. 2022. The role of telomeres in human disease. Annu Rev Genomics Hum Genet. 23:363–381. doi: 10.1146/annurev-genom-010422-091101.

Azad GK, Tomar RS. 2016. The multifunctional transcription factor rap1: A regulator of yeast physiology. Front Biosci (Landmark Ed). 21(5):918–930. doi: 10.2741/4429.

Baur JA et al. 2001. Telomere position effect in human cells. Science. 292(5524):2075–2077. doi: 10.1126/science.1062329.

Brothers M, Rine J. 2022. Distinguishing between recruitment and spread of silent chromatin structures in saccharomyces cerevisiae. Elife. 11. doi: 10.7554/eLife.75653.

Brown CA, Murray AW, Verstrepen KJ. 2010. Rapid expansion and functional divergence of subtelomeric gene families in yeasts. Curr Biol. 20(10):895–903. doi: 10.1016/j.cub.2010.04.027.

Buck SW, Shore D. 1995. Action of a rap1 carboxy-terminal silencing domain reveals an underlying competition between hmr and telomeres in yeast. Genes & Development. 9(3):370–384. doi: 10.1101/gad.9.3.370.

Celli GB, Denchi EL, de Lange T. 2006. Ku70 stimulates fusion of dysfunctional telomeres yet protects chromosome ends from homologous recombination. Nat Cell Biol. 8(8):885–890. doi: 10.1038/ncb1444.

Chatzitheodoridou D et al. 2024. Decoupled transcript and protein concentrations ensure histone homeostasis in different nutrients. EMBO J. 43(21):5141–5168. doi: 10.1038/s44318-024-00227-w.

Chen S et al. 2018. Fastp: An ultra-fast all-in-one fastq preprocessor. Bioinformatics. 34(17):i884–i890. doi: 10.1128/MCB.21.5.1819-1827.2001.

d’Adda di Fagagna F, et al. 2003. A DNA damage checkpoint response in telomere-initiated senescence. Nature. 426(6963):194–198. doi: 10.1038/nature02118.

Destruelle M, Holzer H, Klionsky DJ. 1994. Identification and characterization of a novel yeast gene: The ygp1 gene product is a highly glycosylated secreted protein that is synthesized in response to nutrient limitation. Molecular and Cellular Biology. 14(4):2740–2754. doi: 10.1128/mcb.14.4.2740.

Edgar R, Domrachev M, Lash AE. 2002. Gene expression omnibus: Ncbi gene expression and hybridization array data repository. Nucleic Acids Res. 30(1):207–210. doi: 10.1093/nar/30.1.207.

Ellahi A, Thurtle DM, Rine J. 2015. The chromatin and transcriptional landscape of native saccharomyces cerevisiae telomeres and subtelomeric domains. Genetics. 200(2):505–521. doi: 10.1534/genetics.115.175711.

Gottschling DE et al. 1990. Position effect at s. Cerevisiae telomeres: Reversible repression of pol ii transcription. Cell. 63(4):751–762. doi: 10.1016/0092-8674(90)90141-z.

Harari Y, Zadok-Laviel S, Kupiec M. 2017. Long telomeres do not affect cellular fitness in yeast. mBio. 8(4). doi: 10.1128/mBio.01314-17.

Juarez-Reyes A et al. 2023. Systematic profiling of subtelomeric silencing factors in budding yeast. G3 (Bethesda). 13(10). doi: 10.1093/g3journal/jkad153.

Kalra S et al. 2023. Genome-wide gene expression responses to experimental manipulation of saccharomyces cerevisiae repressor activator protein 1 (rap1) expression level. Genomics. 115(3):110625. doi: 10.1016/j.ygeno.2023.110625.

Keren L et al. 2016. Massively parallel interrogation of the effects of gene expression levels on fitness. Cell. 166(5):1282–1294 e1218. doi: 10.1016/j.cell.2016.07.024.

Kim D et al. 2019. Graph-based genome alignment and genotyping with hisat2 and hisat-genotype. Nat Biotechnol. 37(8):907–915. doi: 10.1038/s41587-019-0201-4.

Kim W, Shay JW. 2018. Long-range telomere regulation of gene expression: Telomere looping and telomere position effect over long distances (tpe-old). Differentiation. 99:1–9. doi: 10.1016/j.diff.2017.11.005.

Kupiec M, Weisman R. 2012. Tor links starvation responses to telomere length maintenance. Cell Cycle. 11(12):2268–2271. doi: 10.4161/cc.20401.

Kyrion G et al. 1993. Rap1 and telomere structure regulate telomere position effects in saccharomyces cerevisiae. Genes Dev. 7(7A):1146–1159. doi: 10.1101/gad.7.7a.1146.

Li B, Lustig AJ. 1996. A novel mechanism for telomere size control in saccharomyces cerevisiae. Genes Dev. 10(11):1310–1326. doi: 10.1101/gad.10.11.1310.

Liao Y, Smyth GK, Shi W. 2014. Featurecounts: An efficient general purpose program for assigning sequence reads to genomic features. Bioinformatics. 30(7):923–930. doi: 10.1093/bioinformatics/btt656.

Love MI, Huber W, Anders S. 2014. Moderated estimation of fold change and dispersion for rna-seq data with deseq2. Genome Biol. 15(12):550. doi: 10.1186/s13059-014-0550-8.

Marcand S et al. 1996. Silencing of genes at nontelomeric sites in yeast is controlled by sequestration of silencing factors at telomeres by rap1 protein. Genes & Development. 10(11):1297–1309. doi: 10.1101/gad.10.11.1297.

McNally EJ, Luncsford PJ, Armanios M. 2019. Long telomeres and cancer risk: The price of cellular immortality. J Clin Invest. 129(9):3474–3481. doi: 10.1172/JCI120851.

Olson CL, Wuttke DS. 2024. Guardians of the genome: How the single-stranded DNA-binding proteins rpa and cst facilitate telomere replication. Biomolecules. 14(3). doi: 10.3390/biom14030263.

Pertea M et al. 2015. Stringtie enables improved reconstruction of a transcriptome from rna-seq reads. Nat Biotechnol. 33(3):290–295. doi: 10.1038/nbt.3122.

Petti AA et al. 2011. Survival of starving yeast is correlated with oxidative stress response and nonrespiratory mitochondrial function. Proc Natl Acad Sci U S A. 108(45):E1089–1098. doi: 10.1073/pnas.1101494108.

Pickett HA et al. 2009. Control of telomere length by a trimming mechanism that involves generation of t-circles. EMBO J. 28(7):799–809. doi: 10.1038/emboj.2009.42.

Rest JS et al. 2013. Nonlinear fitness consequences of variation in expression level of a eukaryotic gene. Mol Biol Evol. 30(2):448–456. doi: 10.1093/molbev/mss248.

Rice AM, McLysaght A. 2017. Dosage-sensitive genes in evolution and disease. BMC Biol. 15(1):78. doi: 10.1186/s12915-017-0418-y.

Rusche LN, Kirchmaier AL, Rine J. 2003. The establishment, inheritance, and function of silenced chromatin in saccharomyces cerevisiae. Annu Rev Biochem. 72:481–516. doi: 10.1146/annurev.biochem.72.121801.161547.

Sidarava V, Mearns S, Lydall D. 2025. Long telomere inheritance through budding yeast sexual cycles. Genetics. 231(1). doi: 10.1093/genetics/iyaf129.

Szafraniec K et al. 2003. Small fitness effects and weak genetic interactions between deleterious mutations in heterozygous loci of the yeast saccharomyces cerevisiae. Genet Res. 82(1):19–31. doi: 10.1017/s001667230300630x.

Teixeira MC et al. 2023. Yeastract+: A portal for the exploitation of global transcription regulation and metabolic model data in yeast biotechnology and pathogenesis. Nucleic Acids Res. 51(D1):D785–D791. doi: 10.1093/nar/gkac1041.

Trapnell C et al. 2010. Transcript assembly and quantification by rna-seq reveals unannotated transcripts and isoform switching during cell differentiation. Nat Biotechnol. 28(5):511–515. doi: 10.1038/nbt.1621.

Ungar L et al. 2011. Tor complex 1 controls telomere length by affecting the level of ku. Curr Biol. 21(24):2115–2120. doi: 10.1016/j.cub.2011.11.024.

Watson JM, Shippen DE. 2007. Telomere rapid deletion regulates telomere length in arabidopsis thaliana. Mol Cell Biol. 27(5):1706–1715. doi: 10.1128/MCB.02059-06.

Wu J et al. 2004. Global analysis of nutrient control of gene expression in saccharomyces cerevisiae during growth and starvation. Proc Natl Acad Sci U S A. 101(9):3148–3153. doi: 10.1073/pnas.0308321100.

Wu L et al. 2006. Pot1 deficiency initiates DNA damage checkpoint activation and aberrant homologous recombination at telomeres. Cell. 126(1):49–62. doi: 10.1016/j.cell.2006.05.037.

Wu T et al. 2021. Clusterprofiler 4.0: A universal enrichment tool for interpreting omics data. Innovation (Camb). 2(3):100141. doi: 10.1016/j.xinn.2021.100141.

Yang D et al. 2011. Human telomeric proteins occupy selective interstitial sites. Cell Res. 21(7):1013–1027. doi: 10.1038/cr.2011.39.

